# Discovery of conformation-sensitive anti-amyloid protofibril monoclonal antibodies using an engineered chaperone-like amyloid-binding protein

**DOI:** 10.1101/558809

**Authors:** W Vallen Graham, Alessandra Bonito-Oliva, Rita Agostinelli, Riyaz Karim, Jeremy Deguzman, Kerry Kelleher, Marianne Petro, Anna-Karin Lindström, Caroline Graff, Kathleen M. Wood, Lioudmila Tchistiakova, Kimberly Marquette, Paul D. Wes, Thomas P. Sakmar

## Abstract

The hypothesis that amyloid beta peptides (Aβ) are central to the pathogenesis of sporadic Alzheimer’s disease (AD) is still hotly debated. Although several monoclonal antibodies (mAbs) against Aβ have failed in therapeutic clinical trials, two conformation-selective, anti-Aβ mAbs continue to show promise. A significant challenge has been to discover mAbs that preferentially target Aβ protofibrils over natively-folded monomeric peptides or amyloid plaques. We have engineered a novel chaperone-like amyloid-binding protein (CLABP), Nucleobindin 1 (NUCB1), which enables the stabilization of protofibrils, allowing them to be used as immunogens in mice to facilitate the generation of mAbs that recognize Aβ protofibrils. An immunization campaign and subsequent screening funnel identified a panel of mAbs with high-affinity to Aβ. Two mAbs in particular, 1A8 and 7C8, displayed significant conformation sensitivity and preferentially bound Aβ protofibrils over monomers. Furthermore, 1A8 delayed Aβ aggregation, but did not prevent eventual fibril formation, while 7C8 significantly and dose-dependently reduced fibril formation by inhibiting both primary and secondary nucleation. Both mAbs protected against protofibril-induced cytotoxicity *in vitro,* and showed distinctive staining patterns by immunohistochemistry in PS1/APP mice and in post-mortem AD brain tissue. In summary, we describe a novel method to stabilize soluble Aβ protofibrils for use in immunization campaigns. We hypothesize that the stabilized protofibrils retain the neoepitopes of the Aβ protofibril and the aggregates found in mouse models of disease and post-mortem AD brain tissue.

## Introduction

While many aging-associated diseases, such as cardiovascular disease, cancer, and hypertension, have benefited from improved diagnostics, medical procedures, and therapeutics, early diagnosis and disease-modifying therapeutics remain urgently needed for Alzheimer’s disease (AD). AD is the leading cause of age-related dementia and is one of the top global healthcare challenges of our time. There are currently 35 million people afflicted with AD and related dementias worldwide, and this number is expected to reach over 100 million by 2050 if prevention strategies are not approved [1, 2].

The clinical manifestations of AD include a progressive increase in confusion and memory deficits, eventually resulting in a diagnosis of cognitive impairment. Pathologically, AD is defined by the presence of amyloid plaques and neurofibrillary tangles, composed of beta-amyloid (Aβ) and tau, respectively. The leading risk factor for AD is age, although genetic and environmental factors contribute to the disease occurrence. The complex multi-domain molecular and cellular pathophysiological nature of suspected disease-causing events has resulted in disagreement in the research community as to the optimal target of potential therapeutics. Currently, there are no approved therapeutics that prevent or even slow the progression of AD or related dementias. As of 2017, there were 54 drugs with disease-modifying indications in Phase II or Phase III clinical trials [3]. Of these entities, 65% target pathways related to Aβ or tau. While many factors are hypothesized to contribute to the etiology of sporadic AD [4, 5], Aβ is likely to play a key role in the molecular pathophysiology of AD. Not only is Aβ the primary component of plaques, but mutations in genes involved in Aβ processing and/or aggregation cause dominant, early-onset familial forms of AD: mutations can be found both in the substrate (amyloid precursor protein, APP) [6] and the protease (presenilin) that work together to form Aβ.

One approach to target and engage Aβ pathology is to use passive immunization with anti-Aβ antibodies. Of the seven immunotherapy drug entities with known anti-Aβ mechanism of action undergoing clinical trials [7, 8], two bind the soluble monomer, four bind to an aggregated form (either oligomers, protofibrils or fibrils), and one is a polyclonal IVIG preparation. While clinical trials for immunotherapies that target the soluble monomer have experienced failures in the past, the emerging approach of targeting soluble Aβ aggregates (oligomers or protofibrils) is showing promising results in ongoing trials [9]. Targeting the soluble pathophysiological form of early Aβ aggregates is appealing, but these conformation-sensitive monoclonal antibodies (mAbs) are extremely hard to discover due to the transient nature and structural polymorphism of the protofibril [10]. The mAb BAN2401 (BioArtic Neuroscience AB, Biogen, Inc., Eisai Co., Ltd.) was developed following the discovery that the Arctic mutation in the APP gene yields a higher propensity for protofibril species compared with fibril formation. Therefore, the protofibrils originating from the Arctic mutant Aβ42 were used to immunize mice and produce a humanized mAb that selectively recognizes protofibrils over monomers and fibrils [11]. Aducanumab (Biogen, Inc.) was discovered using a reverse translational medicine approach whereby B cell libraries were made using blood from aged, cognitively normal human donors. This fully human IgG1 mAb binds to and reduces Aβ aggregates [12].

Stabilizing and maintaining the protofibril structure is challenging. One approach has been to chemically stabilize [13–22] or induce cysteine mutations in the Aβ peptide [23]. However, it is not clear that these Aβ protofibrils result in disease-relevant neoepitopes for mAb production.

An alternative strategy was used to produce the A11 polyclonal antibody [24]. Aβ40 peptide monomers were coated onto gold particles and immunized into mice, yielding a polyclonal pool of antibodies reactive against soluble aggregates of different amyloid sources. An alternative method for Aβ protofibril stabilization is to use naturally existing or engineered amyloid-binding proteins.

Chaperone-like amyloid binding proteins (CLABPs) are an emerging class of proteins that could be useful for capture and stabilization of various amyloid intermediate states. For example, the engineered ZAβ3 affibody binds to and stabilizes the β-sheet hairpin loop in misfolded Aβ monomers that is similar, though not identical, to the β hairpin found in amyloid fibrils [25, 26]. Similarly, the DNAJB6 molecular chaperone binds to misfolded monomers of amyloid from multiple amyloidogenic peptide sources [27–29]. The protein chaperone domain, Bri2-BRICHOS extends the Aβ lag phase of aggregation by maintaining an unstructured monomer [30]. Clusterin, on the other hand, can bind to amyloid oligomers through interaction with biologically active exposed hydrophobic patches [31]. We have recently characterized the ubiquitously expressed protein, Nucleobindin 1 (NUCB1), as a novel CLABP that potently stabilizes amyloid protofibrils from multiple sources [32]. Notably, NUCB1 binding and stabilization preserves the protofibril morphology and detoxifies the amyloid intermediate [32, 33], thereby making the chaperone-like effect of NUCB1 unique in this class of proteins.

We hypothesized that NUCB1-stabilized protofibrils could be isolated and used as immunogens for the discovery of high-affinity, conformation-sensitive mAbs. Here we report the discovery of two such mAbs, 1A8 and 7C8, that preferentially target the protofibril over the monomer, as determined by ELISA and surface plasmon resonance (SPR). These mAbs show functional binding through kinetic inhibition of Aβ42 aggregation and, importantly, display a protective function against Aβ42 in cell toxicity assays. Immunohistochemistry studies on PS1/APP mice showed staining of amyloid plaques, whereas staining of human AD cortex also showed intracellular staining, staining near vessel walls, as well as at dense plaques and diffuse plaque-like structures. Therefore, NUCB1-stabilized Aβ42 protofibrils retain neoepitopes of the protofibril state and can be used to discover high-affinity conformation-sensitive anti-protofibril antibodies.

## Materials and Methods

### Ethics statement

All animal tissue samples were collected from animals in accordance with regulations and established guidelines, including review and approval by Pfizer’s Institutional Animal Care and Use Committee. The Brain Bank at Karolinska Institute provided human tissue from voluntary donations after informed consent. All sections of this report adhere to the ARRIVE Guidelines for reporting animal research [34]. A completed ARRIVE guidelines checklist is included in Checklist S1.

### Production of Aβ42 monomers and protofibrils

Aβ42 and Aβ40 synthetic peptides (American Peptide Company, Cat # 62-0-80) were solubilized in hexafluoroisopropanol (HFIP) at 1 *μ*g/*μ*l for 1 h at room temperature with occasional vortexing, and sonicated in a water bath sonicator (VWR Model 50HT) for 10 minutes. Samples were aliquoted in low-retention tubes (Fisher, Cat # 02-681-320) at 30 *μ*g/tube, dried with a speed vac, and stored at -80 °C. The peptide was then reconstituted in 2 mM NaOH to 1 *μ*g/*μ*l, sonicated in a water bath for 1 minute, dried down in a preheated speed vac for 30 minutes, stored at -80 °C and used within 24 h by diluting in cell media (for the cytotoxicity assay) or 20 mM Sodium Phosphate, pH8.0 (in all other cases). Aβ42 monomers were immediately used for experiments, while Aβ42 protofibrils were obtained by incubating the peptide at 10 *μ*M monomeric concentration for 1 h at 37 °C under quiescent conditions *(i.e.,* without shaking).

### *mt*NUCB1-capped Aβ42 protofibrils

Recombinant expression of the engineered, soluble and Ca^2+^-free sNUCB1 (*mt*NUCB1) has been previously described [32]. 20 *μ*M Aβ42 was co-incubated together with 5 *μ*M *mt*NUCB1 in 20 mM Sodium Phosphate pH 8.0 for 24 h, at 37 °C in quiescent conditions. The capped-protofibril containing solution was then applied to a Superdex200 26/60 PG SEC column (GE Healthcare, Piscataway, NJ) equilibrated with 20 mM sodium phosphate, pH 8.0, 150 mM NaCl. The relevant peak was collected for subsequent experiments.

### Immuno-Electron Microscopy

*mt*NUCB1-capped Aβ42 protofibrils isolated through size exclusion chromatography (SEC) were imaged by double immuno-electron microscopy (EM), as described in [32]. The sample was co-incubated with the mouse anti-Aβ 6E10 (BioLegend, 1:100) antibody and the rabbit anti-NUCB1 (Aviva Systems Biology, 1:100) antibody in solution for 20 min at room temperature. Successively, the sample was diluted to 5 *μ*M and placed in a volume of 5 *μ*l onto a carbon film 200-mesh copper grid for 2 min, followed by a 3 min incubation with 3% BSA. The grid was then incubated for 20 min with an anti-rabbit 12 nm gold-conjugated secondary antibody together with an anti-mouse 6 nm gold-conjugated secondary antibody (Jackson Laboratories, 1:20). The grid was then extensively rinsed in buffer and counterstained with 1% aqueous uranyl acetate solution. Samples were viewed with a JEOL 1400 Plus transmission electron microscope (TEM) and images acquired with Gatan 2K x 2K digital camera.

### Atomic Force Microscopy

The *mt*NUCB1-Aβ42 protofibrils were further imaged by atomic force microscopy (AFM), as previously described [32]. Briefly, the sample was diluted to the desired working concentration and immediately plated (40 *μ*l) on freshly cleaved mica (SPI). After a 10 s incubation, the sample was washed under a gentle stream of 10 ml molecular biology grade H2O (Fisher BP2819-1) before being blown dry with N2 gas and immediately placed under the AFM stage. High-resolution images (1 *μ*m x 1 *μ*m, 512 x 512 pixels) were acquired in air using a combination of the Cypher ES and the MFP-3D-BIO AFMs (Asylum Research, Goleta, CA) and in tapping mode using Olympus AC240TS-R3 probe (Asylum Research, Goleta CA). Each segmented structure was then cropped into its own individual image, a bicubic interpolation was applied, and montages of individual protofibrils were created.

### Immunization campaign

Three BALB/c mice (The Jackson Laboratory) were immunized subcutaneously and intraperitoneally with a total of 20 *μ*g of freshly prepared *mt*NUCB1 -capped Aβ protofibrils, and boosted three additional times every 14 d. Blood was collected 7 d after the second boost. Two mice with high serum reactivity to *mt*NUCB1-Aβ were boosted a final time 2 d after the third boost. 5 d later, the mice were euthanized by low-flow carbon dioxide overexposure followed by exsanguination by cardiac puncture and spleens were harvested for hybridoma fusions.

### Hybridoma fusions and screening

The mouse spleens were fused with P3 myeloma cells (ATCC Cat # CRL1580) according to published methods [35]. Hybridoma supernatants were screened for *mt*NUCB1 and Aβ protofibril reactivity using ELISAs. Of 2,208 supernatants screened, 80 Aβ protofibril-specific hybridomas were selected for retesting using a cut-off value of 0.6 OD. Of 108 *mt*NUCB1-specific hybridomas (OD > 0.6), ten were selected for further testing. In addition, two *mt*NUCB1 and Aβ protofibril cross-reactive hybridomas were selected for further testing. Confirmed hits were further subcloned to ensure monoclonality. Subclones that retained activity were cloned recombinantly into an expression vector containing the human Fc antibody constant region.

### ELISA

In the Sandwich ELISA format, 10 *μ*g/ml of the capture, anti-Aβ N-terminus antibody [36] were coated on clear 96-well plates (Costar #3590) overnight at 4 °C. Aβ42 monomer or protofibril was added at 1 *μ*M and incubated for 1 h at room temperature. Test antibodies were added, incubated for 1 h at room temperature, and successively detected with appropriate HRP conjugated secondary antibodies. ELISAs were developed with TMB One Component HRP Microwell Substrate (TMBW-1000-01) and then stopped with 0.18 M sulfuric acid. Absorbance at 450 nM was read on Envision plate reader (Perkin Elmer).

Similarly, direct ELISA wells were coated with antigen in PBS overnight at 4 °C. Test hybridoma supernatants were added, incubated for 1 h at room temperature, and successively detected with appropriate HRP conjugated secondary antibodies and developed with TMB.

To measure the solution competition of antibodies to immobilized Aβ42 protofibrils by soluble monomeric Aβ40, black 96-well maxisorp plates (NUNC) were coated with Aβ42 protofibrils diluted in coating buffer at a concentration of 2.5 *μ*g/ml (555.6 nM) overnight at 4 °C. The following day the plate was washed and blocked with TBST containing 1% BSA for 1 h at room temperature and successively incubated with a fixed concentration (0.4 nM) of antibody (hu1A8, mt1A8, hu7C8, mt7C8, or the Aβ N-terminus binding 6E10) together with decreasing concentrations of Aβ40 monomers, starting at 10 *μ*M and diluted in half-logs. The binding was detected with appropriate HRP conjugated secondary antibodies and Amplex UltraRed (Thermo # A36006).

### Surface Plasmon Resonance

The surface plasmon resonance (SPR) characterization of an N-terminal Aβ peptide DAE-EG (DAEFRHDSGYSGKQKSRNEGKGGC) binding to anti-mouse captured antibodies from hybridoma supernatants was performed using a BIAcore T-200 instrument (GE Healthcare, Marlborough, MA) [36]. The sample and running buffer was HBS-EP, pH 7.4 (10 mM HEPES, 150 mM NaCl, 3 mM EDTA, and 0.05% P20). An anti-mouse antibody (GE Healthcare, BR100838) was covalently coupled to a CM5 sensor chip (GE Healthcare, BR100530) following the manufacturer’s recommendations. Mouse hybridoma antibodies at 0.5 *μ*g/mL were captured for 120 s on flow cells 2, 3, or 4 with flow cell 1 used as a reference. The DAE-EG peptide was diluted to 500 nM and injected over the sensor chip surface for 120 s at a flow rate of 50 *μ*l/min. Dissociation data was collected for 300 s post injection followed with three 60 s regeneration pulses of 10mM Glycine pH 1.5, and 1 pulse of HBS-EP, pH 7.4. The sensorgram data was collected at 1Hz, double referenced [37], and fit to a 1:1 Langmuir model using BIAcore T200 evaluation software version 3.0. The triage quality equilibrium dissociation constant KD was determined with the equation K_D_ = kd (1/s) / ka (1/Ms).

The SPR studies to determine conformation binding of the human chimeric 1A8 and 7C8 mAbs were carried out with the ProteOn XPR36 protein interaction array system (Bio-Rad) based on SPR technology. The antibodies (1A8 and 7C8) were immobilized in the vertical direction on GLM sensor chips (Bio-Rad) using amine-coupling chemistry, as described previously [38], followed by a blocking step with ethanolamine. The anti-Aβ 6E10 (Bio-Legend) antibody and the mouse IgG antibody 1D4 were used as positive and negative control, respectively. The final immobilization level was about 6500 resonance units (1 resonance unit = 1 pg protein/mm^2^) for all the antibodies. Successively, Aβ42 protofibrils obtained by incubating 10 *μ*M Aβ42 for 60 min at 37 °C or freshly solubilized Aβ40 monomers were diluted in 20 mM Sodium Phosphate pH 8.0 and flowed over the chip surface, in the horizontal direction, for 60 s at a flow rate of 30 *μ*l/ml. The assays were performed at 25 °C and the data were normalized by interspot and by buffer.

### Octet

An OctetRED 384 instrument (ForteBio, Menlo Park, CA) was used to characterize binding of *mt*NUCB1-capped Aβ42 protofibrils and freshly prepared protofibrils without *mt*NUCB1 to mouse hybridoma antibodies. The mouse hybridoma antibodies were diluted to 10 *μ*g/ml in PBS and loaded for 400 s onto an anti-mouse Fc biosensor (ForteBio, 18-5089). A baseline was established in Kinetics buffer (ForteBio, 18-5032) for 180 s, followed by a 300 s association and dissociation of *mt*NUCB1-Aβ42 at ~300 nM or protofibrils at ~900 nM. The Octet assay was conducted at ambient temperature. The data was double referenced [37] and analyzed with BIAevaluation software version 4.1.1.

### Fab fragments purification

1A8 and 7C8 IgGs, as well as the control IgG, were digested with immobilized papain protease and Fab fragments were purified using Protein A agarose (Pierce Fab Preparation Kits). The success of the digestion reaction was assessed by native SDS-PAGE gel.

### Thioflavin T binding assay

The kinetics of aggregation of Aβ40 (40*μ*M) was measured by incubating the peptide with thioflavin T (ThT) (10 *μ*M) (Fisher Scientific). The kinetics of aggregation of Aβ42 (10 *μ*M) was tested in the presence of equimolar concentration of the whole IgG antibody 1A8, 7C8, or negative control (10 *μ*M, Ultra-LEAF Purified Human IgG1 isotype control, Biolegend #403502) or different concentrations (2.5, 5, 7.5 or 10 *μ*M) of their corresponding digested Fab fragments and ThT (10 *μ*M). A volume of 50 *μ*l per well (n = 4/group) was added to each well of a prechilled (4 °C) Corning 96-well half area black with clear flat bottom polystyrene with non-binding surface (NBS) and covered with clear self-adhesive topseal. The aggregation was tested every 10 min under quiescent conditions for up to 7 d at a constant temperature of 25 °C (for Aβ40), or 24 h at 37 °C (for Aβ42). Fluorescence measurements were performed on a Flexstation II (Molecular Devices) using an excitation wavelength of 450 nm and an emission wavelength of 485 nm. The obtained fluorescence measures were normalized to the relative fluorescence expressed after 20 min of incubation.

### Rate constant calculation

Rate constants of aggregation (primary nucleation, elongation and secondary nucleation) were calculated with AmyloFit online software [39] and simulations were performed to determine how antibody Fab affects the global Aβ42 aggregation profile by interfering with and inhibiting microscopic aggregation event(s). First, following the preliminary steps in the AmyloFit pipeline, we chose time windows from reaction start point to plateau and normalized the values to 1. The fitted model of secondary nucleation dominant aggregation was chosen according to the guidelines published in [40]. Successively, the model parameters [initial monomer concentration (m0), initial fibril number concentration (P0), initial fibril mass concentration (M0), reaction order of primary nucleation (nc) and reaction order of secondary nucleation (n2)] were set to Global constant (see tables for specific values) and each time one of the rate constants was set to ‘Fit’ while the others were set to ‘Global fit’. We made sure convergence was attained by increasing the Basin Hops and observing no change in the MRE. The fitting results expressed as MRE and residuals over reaction time were analyzed and shown in each of these specific fittings separately.

### Cytotoxicity assay

Cytotoxicity assays were performed using adherent PC12 cells (CRL01721.2, ATCC). Experiments were only conducted if cells were above 90% viable. Cells were diluted to a concentration of 3.2 x 10^5^ cells/ml in assay media: DMEM/F-12, no phenol red (GIBCO, Cat # 21040-025), 0.5% FBS, non-heat inactivated (ATCC, Cat # 30-2020). 80 *μ*l of cell suspensions were added to flat-bottomed, black-walled 96-well plates (Corning, Cat # 3340 CellBIND), for a final amount of 25,600 cells per well, and incubated overnight. On the following day, NaOH-treated Aβ42 was diluted in assay media, sonicated in a water bath for 1 min and either used immediately to test monomerics, or incubated at 10 *μ*M monomeric concentration at 37 °C for 1 h, protected from light, to test protofibrils, and successively diluted to the final concentration.

To determine the EC80 of monomers and protofibrils, the samples were added to cells seeded the day before at an initial concentration of 1 *μ*M and 3 *μ*M, respectively, diluted half logs across the plate and incubated overnight.

To test the protective effect of the antibodies, 20 *μ*l of Aβ42 monomers or protofibrils were added at EC80 concentrations (800 nM or 30 nM, respectively) to cells seeded the day before and the concentration kept fixed across the plates. At the same time, 1A8, 7C8, or negative control were added to the peptide at over a range of concentrations. Cells were incubated overnight, and viability was measured using the Roche MTT (3-(4,5-dimethylthiazol-2-yl)-2,5-diphenyl tetrazolium bromide) kit (Roche, Cat # 11465007001) according to manufacturer’s instructions.

### Immunohistochemistry in mouse tissue

Adult (male) PS1(G384A)/APPsw (PS1/APP) transgenic mice were ordered from Taconic (Pfizer generated line), housed in groups of four and given 5 d to acclimate to the housing facility. Environmental conditions were compliant with approved protocols. During housing, animals were monitored twice daily for health status. No adverse events were observed.

PS1/APP transgenic mice and non-transgenic littermate controls were euthanized by low-flow carbon dioxide overexposure followed by exsanguination by cardiac puncture, and perfused with saline. Brains were harvested, flash frozen in isopentane, sliced at 12 *μ*m thickness and stored at -80 °C. At the time of use, sections were thawed for 5 min at room temperature. Microscope slides were submerged in Coplin jars containing 4% paraformaldehyde for 60 min to prepare post-fixed sections for staining. Antigen retrieval was performed using Rodent Decloaker (Biocare Medical, Walnut Creek, CA, Cat # RD913) solution in a Decloaking Chamber (Biocare Medical, Walnut Creek, CA, Cat # DC2012). To inactivate endogenous peroxidases and block non-specific binding sites, sections were treated with 0.3% hydrogen peroxidase, an Avidin/Biotin Blocking Kit (Vector Laboratories, Burlingame, CA, Cat # SP-2001), and 10% normal goat serum (Vector Laboratories, Burlingame, CA, Cat # S-4000). Primary antibodies 1A8, 7C8, a total anti-Aβ mAb and isotype control were prepared at 10 *μ*g/ml and incubated with sections overnight in a humid chamber at 4 °C. The following day, sections were incubated with biotinylated secondary antibody (Bethyl Laboratories, Montgomery, TX) for 60 min. To visualize staining, sections were treated with the VECTASTAIN ELITE ABC-HRP Kit (Vector Laboratories, Cat # PK-6100) and DAKO DAB+ Chromogen System (Agilent, Santa Clara, CA, Cat # K3468) per manufacturer protocols. Slides were counterstained with hematoxylin, mounted, and then imaged on the Axio Scan.Z1 slide scanner (Carl Zeiss Microscopy GmbH, Jena, Germany).

### Immunohistochemistry in human tissue

All brain materials were obtained from the Huddinge Brain Bank at Karolinska Institutet Alzheimer Disease Research Center. All familial AD subjects met the criteria for definitive AD according to the Consortium to Establish a Registry for AD (CERAD) 38. 1A8 and 7C8 target engagement was evaluated in formalin-fixed paraffin-embedded (FFPE) tissue sections from the frontal cortex of a patient diagnosed with definitive AD by CERAD and Braak V-VI as well as cerebral amyloid angiopathy. 5 *μ*m sections were sliced and stained with the murine version of the mAbs (1A8 muIgG2a-4m and 7C8 muIgG2a-4m) and compared to the anti-Aβ N-terminus positive control antibody (6E10, Biolegend, Cat. # #SIG39320, diluted 1:1000) and an IgG negative control antibody (used at a concentration of 5 *μ*g/ml). The FFPE sections were deparaffinized in xylene and rehydrated in decreasing concentrations of ethanol. To retrieve antigens, all slides were treated with Diva Decloaker (Biocare Medical, Cat. # DV2004MX) in a pressure cooker at 110 °C for 30 min with the exception of the 6E10 stain slide that was treated with 70% Formic Acid for 20 min at room temperature. Endogenous peroxidase activity was successively blocked with Peroxidase (Dako kit, Cat. # K4007) and the sections were incubated with normal goat serum diluted 1:20 in TBST for 20 min at room temperature. The sections were then incubated for 45 min room temperature in the respective primary antibody diluted in antibody diluent (Dako, Cat. # S3022) and the signal detected using a HRP-DAB based detection system kit (Dako, Cat. # K4007). Finally, nuclei were counterstained with Mayer’s Hematoxylin using a standard protocol. After that, the sections were dehydrated in rising ethanol concentrations followed by clearing in xylene. The images were acquired with a Nikon Eclipse E800 microscope equipped with a 10x objective and the NIS-Elements F 4.30.01 software. Human sample collection and the protocols used in the study were approved by the Stockholm ethical review board, unit 1 (Stockholms regional etikprövningsnämnd avdelning 1) with the reference number 2011/962-13/1 on July 20, 2011 and all methods were performed in accordance with the relevant guidelines and regulation thereby established. The tissue was collected post-mortem at the Brain Bank at Karolinska Institute upon voluntary donation and informed consent (informed consent forms are available upon request).

### Statistical analysis

Data were expressed as means ± standard error (SEM). One- and Two-way analysis of variance (ANOVA) followed by a post-hoc Tukey’s multiple comparisons test was used to analyze differences among groups. Statistical analyses were performed using GraphPad Prism 6.0 software (GraphPad Software Inc). Statistical differences for all tests were considered significant at the p<0.05 level.

## Results

### Preparation and characterization of stabilized Aβ42 protofibril immunogen

Previously, we characterized an engineered version of the CLABP, *mt*NUCB1, as an inhibitor of amyloid aggregation, particularly Aβ42 [32]. The conversion of Aβ42 monomers into fibrils is incomplete in the presence of *mt*NUCB1 through a proposed mechanism of *mt*NUCB1 capping Aβ42 protofibril ends. Importantly, *mt*NUCB1 maintains Aβ protofibril solubility and renders the aggregates non-toxic in a cell-based assay. The stable *mt*NUCB1-Aβ42 structures display a characteristic morphology of short protofibrils, being approximately 30 nm long, and 3.8 nm thick, and can be purified using size-exclusion chromatography (SEC) [32]. We hypothesize here that mŕNUCB1-stabilized Aβ42 protofibrils contain an array of neoepitopes that characterize the protofibril structure. Therefore, we aimed to discover high-affinity mAbs that bound to the quaternary conformations of the Aβ protofibril. To this end, we purified sufficient *mt*NUCB1-Aβ42 protofibril complex to complete an immunization campaign of three mice.

We generated and enriched *mt*NUCB1-Aβ42 stabilized protofibrils using SEC and the relevant peak was collected and concentrated, as previously reported [32]. The Aβ and NUCB1 content was confirmed with direct ELISA (S1 Fig A, B). Atomic force microscopy (AFM) was utilized to validate that the *mt*NUCB1-Aβ42 complex retained the expected morphology as seen before, with similar length and height as previously observed [32] (S1 Fig C). Immuno-electron microscopy (EM) analysis of NUCB1- and Aβ-directed gold particles showed close proximity to an apparent protofibril structure (S1 Fig D). We therefore considered the immunogen to be robust and representative of *mt*NUCB1-Aβ42 protofibril complexes.

### Screening strategy and summary of screening campaign results

The screening strategy and a summary of screening campaign results are illustrated in the schematic in Fig 1. The hybridoma supernatant fractions resulting from the immunization campaign were compared with pre-fusion bleeds, or negative controls, in an ELISA to detect mŕNUCB1-specific and Aβ42-protofibril-specific activities (Fig 1 and Fig 2A). Relative binding activities of the samples indicate ranges of activity for Aβ42 protofibrils and *mt*NUCB1 separately, and a few supernatant fractions from the hybridoma wells that appear to have dual activity at varying levels, possibly due to mixed hybridoma populations. From the primary screening, a total of 80 hybridomas having Aβ42 protofibril-binding activity over 0.6 OD, and *mt*NUCB1 specific activity below 0.6 OD, were selected for further characterization (Fig 2A, blue circles). Of these, 57 fused lines were confirmed to be monoclonal and re-screened by single-point ELISA to compare binding to Aβ42 monomers, protofibrils, *mt*NUCB1, and the original immunogen *mt*NUCB1-Aβ42 protofibrils (Fig 2B). The results in this single-point ELISA show that many clones displayed preliminary desired characteristics of higher relative affinity to Aβ42 protofibrils than *mt*NUCB1. Some clones showed reactivity both to *mt*NUCB1 -Aβ protofibrils and *mt*NUCB1, suggesting that they may be reactive to *mt*NUCB1.

**Figure 1.**
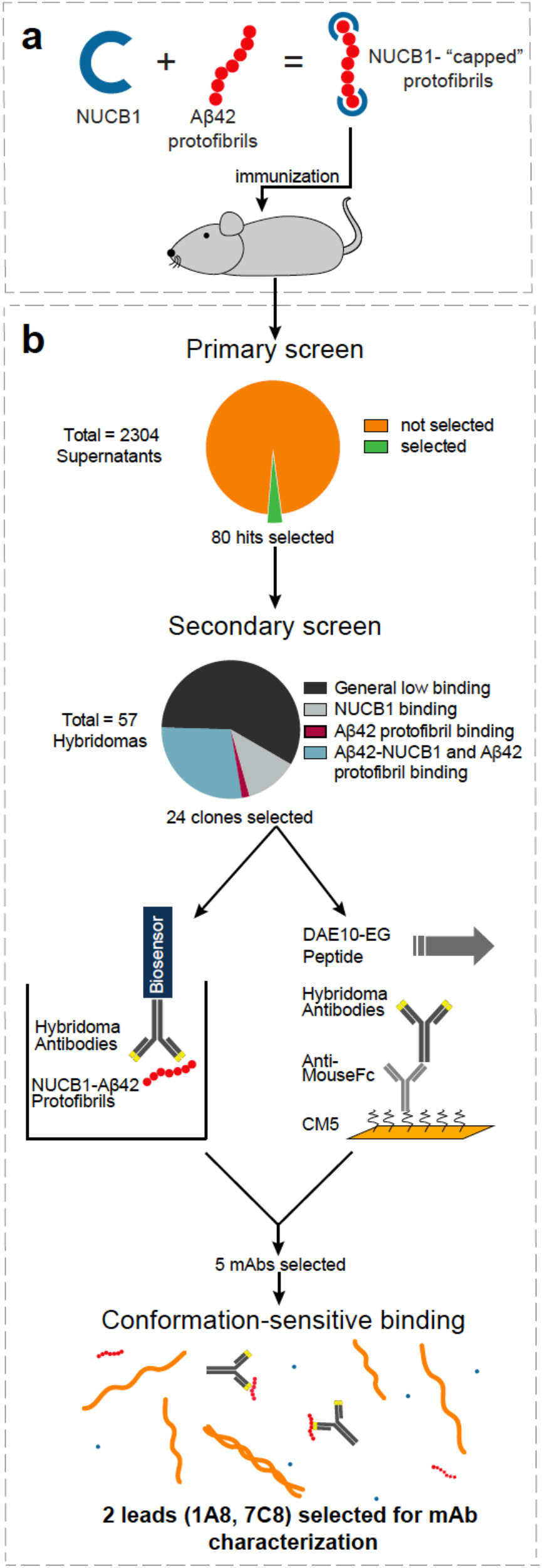
Screening strategy flow chart and immunization campaign summary results. (A) Work flow schematic of the immunogen preparation. (B) Screening strategy flow chart. The hybridoma supernatants (n = 2,304) were subjected to the primary screening by ELISA. Based on reactivity, 80 hits were selected and fused, and 57 clones were submitted to secondary screening by single-point ELISA. Twenty-four supernatants were subsequently tested by Octet Biosensor and Surface plasmon resonance (SPR) assays. Five selected clones were subcloned and screened for protofibril binding by sandwich ELISA. Finally, 2 lead mAbs were selected for further characterization.

**Figure 2.**
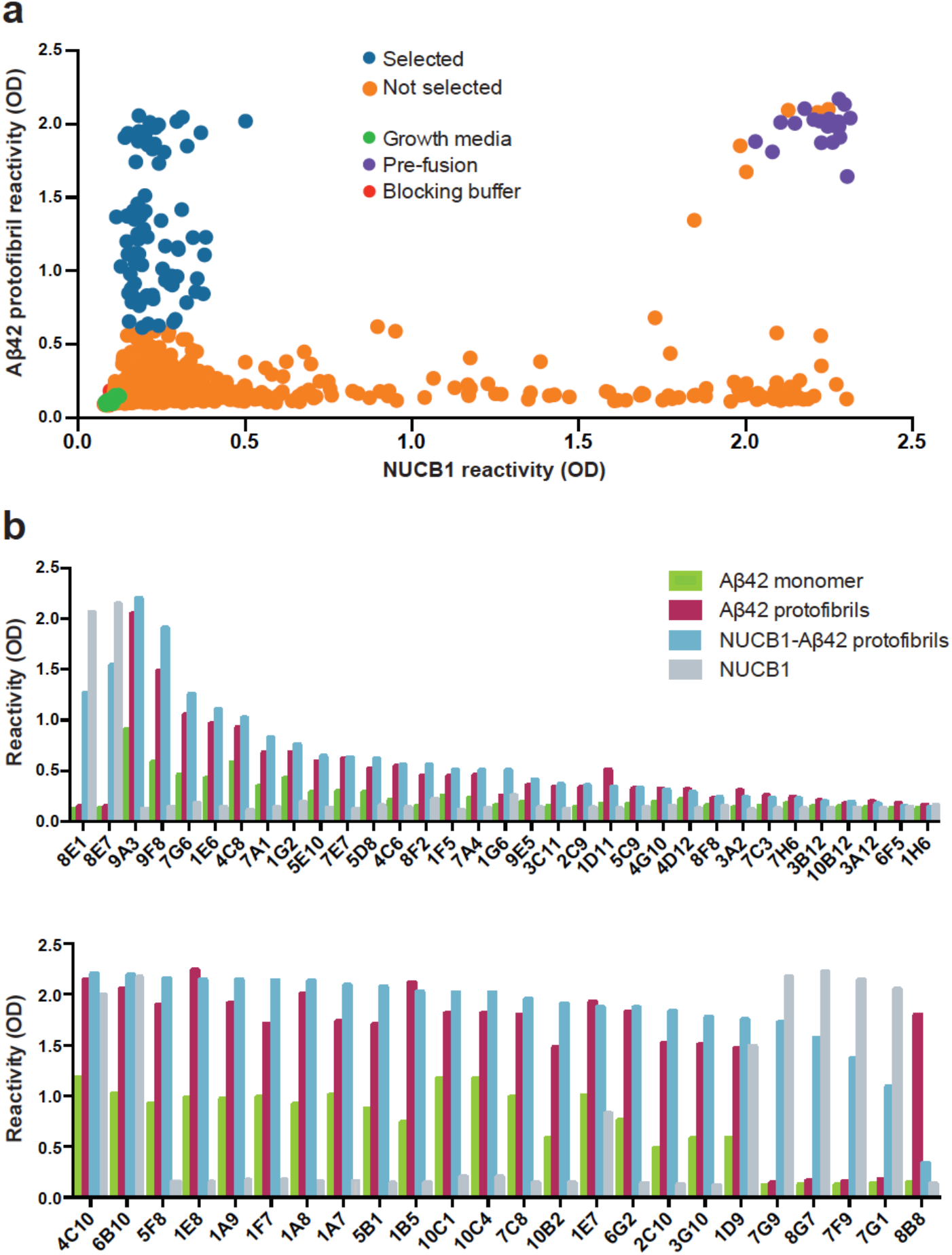
Generation of mAbs recognizing mtNUCB1-capped Aβ protofibrils. (A) Hybridoma cells were generated from the splenocytes of immunized mice and plated at 20,000 cells per well in 96-well plates. The supernatants of 2,304 hybridomas were screened for reactivity against Aβ protofibrils and mtNUCB1. Hits that showed reactivity > 0.6 OD at 450 nm for Aβ protofibrils, but < 0.6 for mtNUCB1, were selected. (B) 57 clones were re-tested for reactivity to mtNUCB1-Aβ protofibrils and mtNUCB1, as well as uncapped Aβ protofibrils and Aβ monomers, in a single-point ELISA. Based on the reactivity profile, 24 hybridomas (shown in the lower panel) were selected for further analysis.

Based on their binding profile in this secondary screening, we selected 24 hybridomas (shown in Fig 2, lower panel) for further binding analysis. Specifically, of these clones, 16 showed high binding to Aβ42 protofibrils and mtNUCB1-Aβ protofibrils, seven clones showed high *mt*NUCB1 binding and one clone displayed preferential binding to Aβ42 protofibrils.

### Octet Characterization of Hybridoma Supernatants

To further determine the relative reactivity of 24 selected hybridoma supernatant antibodies against the immunogen (*mt*NUCB1-Aβ42 protofibrils) or Aβ42 protofibrils, we utilized a label free high-throughput Octet assay (Fig 1 and S2 Fig A). The results indicated that about half of the clones retained their antibody binding activity to both the immunogen and the Aβ42 protofibrils. While some supernatant antibodies lost their binding activity to both antigens *(i.e.,* 1A7, 3G10, 10B2, 10C4, 10B12, 1E8 and 8B8), a few *(i.e.,* 4C10, 7F9, 8G7 and 7G1) showed strong binding to the immunogen, but lost activity against the Aβ protofibrils.

### SPR Characterization of Hybridoma Supernatant Antibodies

An SPR assay was used to characterize binding of the selected hybridoma supernatant antibodies to the N-terminal, immune-dominant region of Aβ [41]. To determine relative binding, a peptide containing the N-terminal 1-10 amino acids of Aβ was screened against the hybridoma supernatant antibodies (Fig 1). The results indicated that half of the supernatant antibodies bound to the N-terminus Aβ 1-10 peptide (S2 Fig B). We selected a total of five supernatants that demonstrated binding to the N-terminus Aβ 1-10 peptide *(i.e.,* 1A8, 7C8, 1D9, 1E7 and 6G2), with Kd values near 10 nM, including 1A8 and 7C8. 1A8 and 7C8 showed the lowest and highest relative binding signal, respectively, though this difference may have been driven by different concentrations and degrees of aggregation in the relatively crude supernatants. All selected clones were processed for sequencing, subcloning and further characterization.

### Monomer versus protofibril binding affinities of cloned mAbs

Antibody was purified from the five selected clones and tested against Aβ42 monomers and protofibrils using a sandwich ELISA (S3 Fig). The range of EC50s of each mAb for Aβ42 monomers ranged from 2.1 to 9.2 nM, while the range for protofibrils was from 2.8 to 5.8 nM (S3 Fig). While the EC50s are in close range, the overall signal for protofibril was higher than monomer for all antibodies, thus indicating that these antibodies have increased binding to the protofibril preparation. These results indicate that the *mt*NUCB1-Aβ42 protofibril complex contains epitopes that are immunogenic, resulting in mAbs that have affinity to Aβ.

### Conformation-sensitive binding of 1A8 and 7C8 expressed mAbs

To produce stable antibody chimeras, we cloned the variable regions of both 1A8 and 7C8 and incorporated these sequences onto a human IgG1 antibody backbone, with or without effector function, in suitable vectors for expression in mammalian cell lines. The effectorless mutant contains point mutations (L234A, L235A, and G237A) in the antibody heavy chain to decrease Fc receptor binding [42, 43]. To validate that these clones retained their desired activity, we purified the human chimeric (hu1A8 and hu7C8) and the effectorless mutant (mt1A8 and mt7C8) mAbs and used them in a sandwich ELISA (Fig 3A, B) to compare relative binding to either Aβ42 protofibrils or freshly prepared Aβ40 monomer. In these experiments the Aβ40 peptide was used as monomeric control because of its slow aggregation rate (S4 Fig), since Aß42 monomer would begin to aggregate during the course of the assay. The effectorless mAbs bound equivalently to their wild-type counterparts, with more binding occurring to lower concentrations of protofibrils, and achieving higher saturable binding than monomers. Since the apparent preferential binding of immobilized 1A8 and 7C8 to protofibrils in the ELISA format could be due to non-physiological avidity effects, we sought to determine whether preferential binding was also observed in a solution phase. Therefore, we performed an ELISA-based solution competition assay, which was recently used to demonstrate preferential binding of aducanumab to Aβ oligomers (44). While monomeric Aβ40 was able to compete for binding to the total Aβ N-terminal mAb, 6E10, with an EC50 of 95nM, minimal competition was observed for 1A8 and 7C8, with only about 30% inhibition observed at the highest concentration tested (3 *μ*M) (Fig 3C, D). The results indicate that our mAbs, although recognizing the Aβ N-terminus, are conformation-sensitive and preferentially bind to the protofibrils even in presence of excessive concentrations of monomers, contrary to the positive control Aβ N-terminus antibody 6E10.

**Figure 3.**
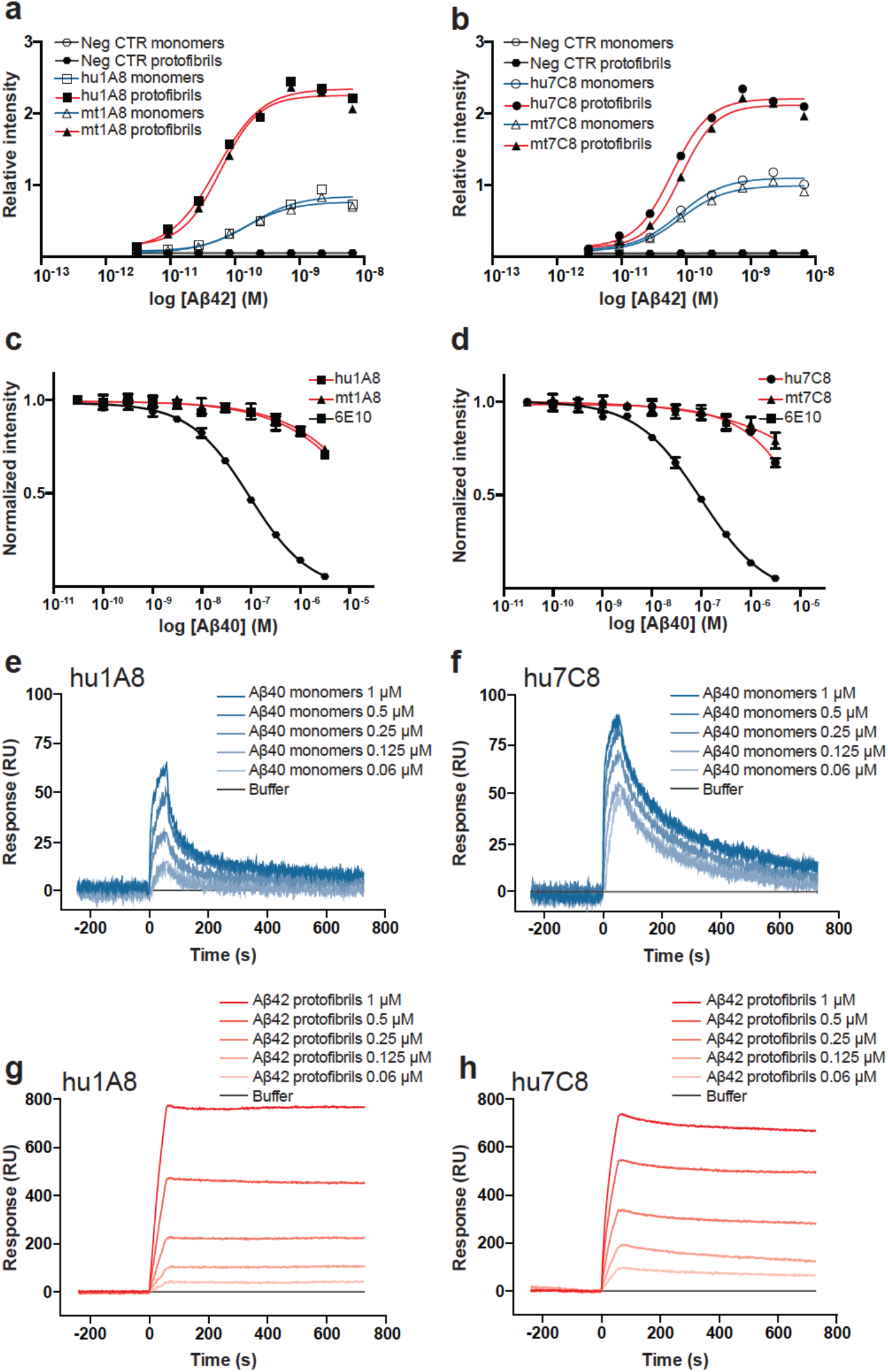
Conformation-sensitive binding of 1A8 and 7C8 to Aβ42 monomers and protofibrils. The relative binding of 1A8 and 7C8 to Aβ42 monomers and protofibrils was tested by (A-D) ELISA and (E-H) surface plasmon resonance (SPR). (A,B) Sandwich ELISA where human chimeric mAbs hu1A8 and hu7C8 (wildtype or effectorless mutant (mt)) are the capture antibodies for either Aβ42 protofibrils or soluble monomeric Aβ40. (C, D) Solution competition of antibodies (hu1A8, mt1A8, hu7C8, and mt7C8) to immobilized Aβ42 protofibrils by soluble monomeric Aβ40, compared to the positive control 6E10 antibody. For the competition ELISA, Aβ42 protofibrils were coated on the ELISA plate and incubated with the antibodies together with decreasing concentrations of Aβ40 monomers. The binding of (E,G) hu1A8 and (F,H) hu7C8 to (E,F) Aβ40 monomers and (G,H) Aβ42 protofibrils was tested by SPR. Aβ40 monomers and Aβ42 protofibrils were prepared at a 10*μ*M initial concentration, diluted to 1, 0.5, 0.250, 0.125, and 0.06*μ*M and flown for 60 s over each antibody (300 nM) previously immobilized on the chip (RL = 6500). Data are normalized by interspot and buffer and presented as mean ± standard error of the mean (SEM).

Conformation-sensitivity was further characterized with a label-free SPR binding assay comparing monomer and protofibril affinity. The assay was validated using either the N-terminal binding anti-Aβ mAb, 6E10 (S5 Fig A, C), or a negative control mAb (S5 Fig B, D). The hu1A8 (Fig 3E) and hu7C8 (Fig 3F) mAbs immobilized on the chip surface showed a mild concentration-dependent binding to Aβ40 monomers (Fig 3E, F), characterized by a rapid dissociation phase. In contrast, the antibodies binding to Aβ42 protofibrils (Fig 3G, H) resulted in a potent and concentration-dependent association curve and lack of dissociation in the 800 seconds of observation.

### Functional inhibition of Aβ42 aggregation by 1A8 and 7C8

Proteins that bind intermediate amyloid aggregates can have an effect on microscopic rate constants of aggregation kinetics [39]. Since 1A8 and 7C8 have high affinity to the intermediate species (protofibrils), we aimed to determine the effect on microscopic aggregation events in the kinetic reaction. The whole molecule mAbs show inhibitory effect on equimolar concentration of Aβ42 aggregation, as measured by the fluorescent signal decrease obtained with 1A8 and 7C8 in a thioflavin T (ThT) kinetic reaction, compared with the control antibody (Fig 4A) (two-way ANOVA with repeated measure indicates a significant time * treatment interaction F_(300,755)_ = 34.23, p < 0.0001, time effect F_(146,2774)_ = 457.0, p < 0.0001 and treatment effect F_(6,19)_ = 91.77, p < 0.0001). However, whereas 7C8 completely inhibits aggregation, 1A8 delays the initial reaction, although it fails to reduce final fibril content (Fig 4A) (one-way ANOVA on the endpoint fluorescence indicates significant treatment effect: F_(2,5)_ = 70.59, p < 0.01, followed by Dunnett’s multiple comparison test).

**Figure 4.**
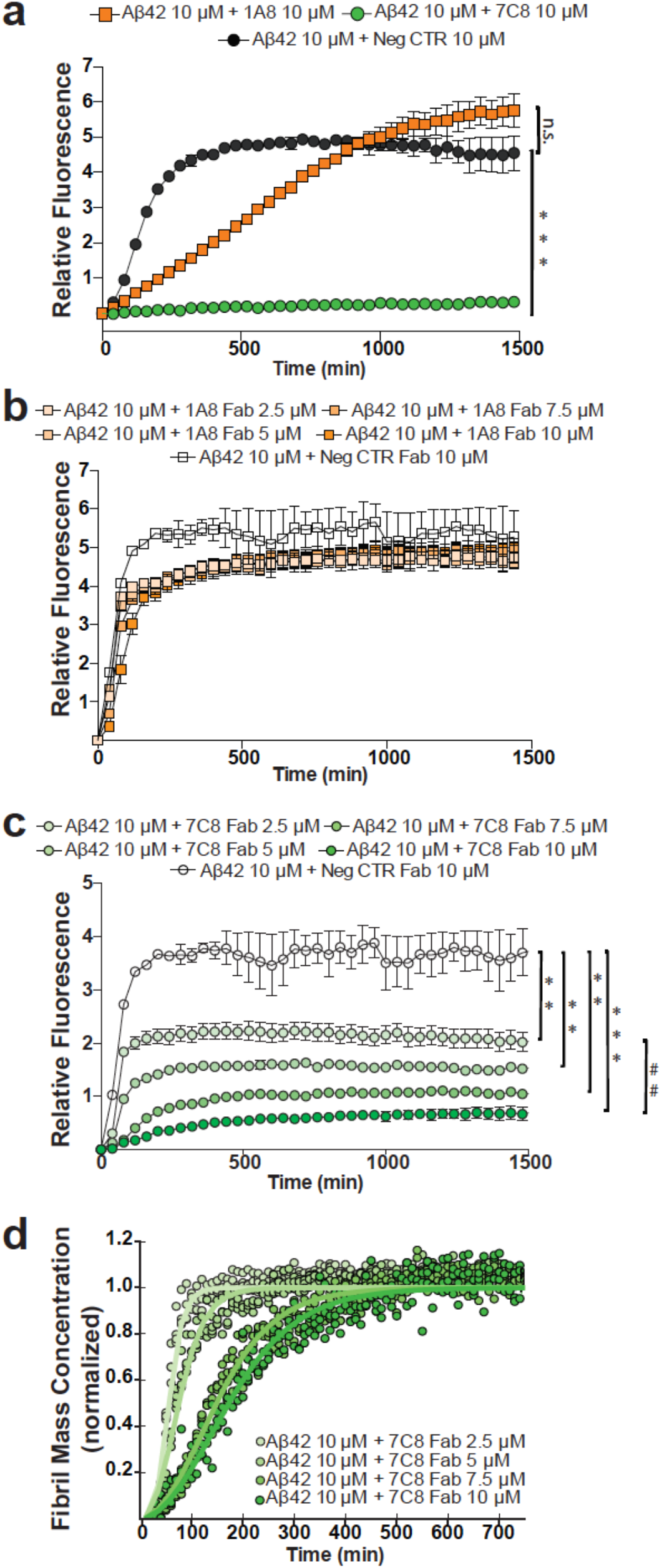
Inhibition of Aß42 aggregation by 1A8 and 7C8. (A) Kinetics of aggregation of 10μM Aß42 incubated at 37 °C for 24 h in quiescent conditions, in presence of 10 μM ThT and equimolar concentrations of 1A8 and 7C8 IgG, increasing concentrations (2.5, 5, 7.5 and 10μM) of digested (B) 1A8 and (C) 7C8 Fab fragments. An IgG1 isotype control and the correspondent digested Fab fragments were used as negative controls. \*\**p*<0.01, \*\*\**p*< 0.0001 vs CTR antibody; ##*p*< 0.01 vs lower antibody concentration (post-hoc comparison). (D) The aggregation kinetics of Aß42 in presence of increasing concentrations (2.5, 5, 7.5 and 10 μM) of digested 7C8 Fab fragments (shown in panel C) was analyzed by Amylofit. For each mAb concentration the aggregation values were normalized to 1 and the rate constant parameters for secondary nucleation were individually fitted in parallel to global fitting of primary nucleation and elongation. Fitted (lines) and the experimental (circles) data are shown.

Subsequently, we obtained digested Fab fragments in order to control the antibody avidity effect and reduce the size of the inhibitory molecule. We found that 1A8 Fab did not show significant functional effect when tested for the inhibition of Aβ42 aggregation in the ThT assay (Fig 4B). The statistical analysis (two-way ANOVA with repeated measure) indicates a significant time * treatment interaction (F_(584, 1898)_ = 6.058, p < 0.0001) and time effect (F_(146, 1898)_ = 488.0 p < 0.0001), but no significant treatment effect.

On the other hand, the 7C8 Fab fragments retain the inhibitory effect expressed by the full 7C8 IgG and potently and concentration-dependently decrease Aβ42 aggregation and final fibril mass (Fig 4C). The inhibitory effect is expressed by both a delay in the aggregation, as seen by the progressive rightward shift of the curves with increasing Fab fragment concentration, and a significant reduction in the endpoint fluorescence, indicating decreased fibril mass after 24h of co-incubation. The statistical analysis of the time course aggregation kinetics revealed a significant time * treatment interaction [two-way ANOVA with repeated measure (time) F_(920, 2990)_ = 10.33 p < 0.0001, time effect F_(230, 2990)_ = 137.6 p < 0.0001 and treatment effect F_(4, 13)_ = 75.93 p < 0.0001], and the analysis of the final fluorescence level indicates a significant treatment effect [one-way ANOVA F_(4, 13)_ = 15.87 p < 0.0001, followed by Tukey’s multiple comparisons test].

Recently, the rate kinetics for Aβ aggregation have been scrutinized, allowing for the understanding of inhibitory effects on each microscopic mechanism [39]. Using the Amylofit software [40], we applied the curve fitting parameters to our primary data of 7C8 Fab-mediated Aβ aggregation and found that the binding domain of the antibody inhibits aggregation with a dual mechanism (Fig 4D). The rate kinetics best fit with a multi-step secondary nucleation dominated model, thereby secondary nucleation as well as primary nucleation kinetics are affected by 7C8 Fab fragments (Fig 4D, S1 Table).

### Rescue of Aβ42-induced cell toxicity by 1A8 and 7C8

To determine whether 1A8 and 7C8 also bound to Aβ in a biological context, we tested whether these antibodies could rescue Aβ-induced cytotoxicity in a cell-based system. To establish the assay, we treated adherent PC12 cells with a concentration range of either Aβ42 monomers or protofibrils, and measured viability using the MTT reagent (Fig 5A). While we do not believe this assay is of physiological relevance to AD *per se* [44, 45], it provides a facile screen to measure binding of the antibodies to Aβ in a biological matrix. The addition of Aβ42 protofibrils was far more toxic than that of monomers, with EC50 around 10 nM compared to 200 nM (Fig 5A). We next treated the cells with 1A8 and 7C8 at the EC80 concentration of either Aβ42 monomer (Fig 5B, D) or protofibril (Fig 5C, E) (30 nM and 800 nM, respectively). Both 1A8 and 7C8 were able to rescue Aβ42-mediated toxicity with nanomolar potencies, though 1A8 antibody variants had a reduced maximal effect compared with 7C8 variants.

**Figure 5.**
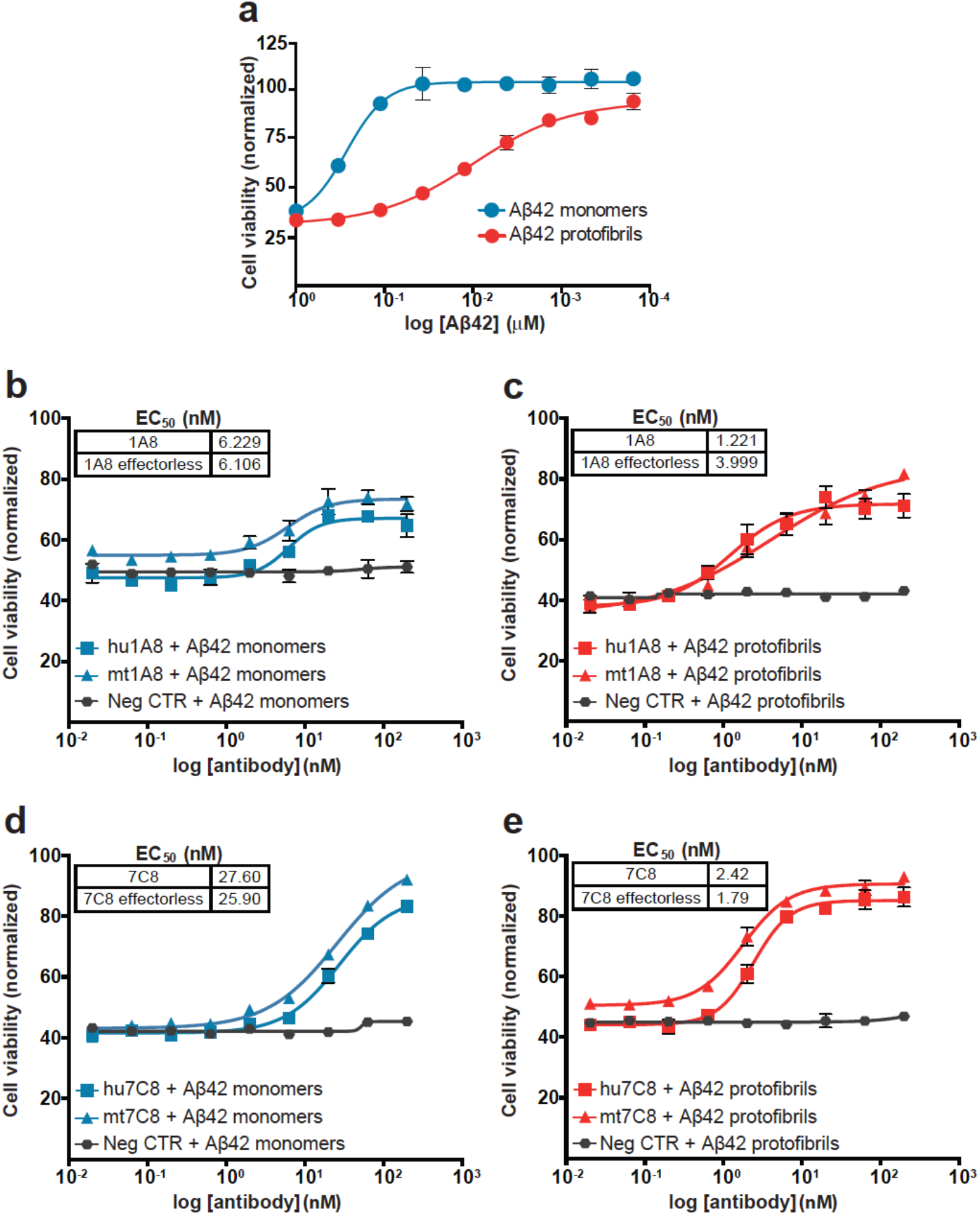
Inhibition of Aβ42-mediated cell toxicity by 1A8 and 7C8. A) Cytotoxicity mediated by Aβ42 monomers and protofibrils in PC12 cells. The viability of adherent PC12 cells was measured by MTT assay in presence of increasing concentrations of Aβ42 monomers or protofibrils, added to the plate at an initial concentration of 1 μM and 3 μM, respectively, and incubated overnight. (B,D) The viability of adherent PC12 cells was measured by MTT assay in presence of Aβ42 monomers (800 nM) or (B,D) protofibrils (30 nM) and increasing concentrations of the human chimeric (B,C) 1A8 (hu1A8) or (D,E) 7C8 (hu7C8) and their respective effectorless proteins (mt1A8 and mt7C8). In each experiment, a human IgG1 was used as a negative isotype control, Aβ42 plus a concentration range of antibodies was incubated with the cells overnight. Viability was then measured using the MTT kit.

### 1A8 and 7C8 staining of PS1/APP mouse brain

To determine if 1A8 and 7C8 could stain amyloid aggregates in tissue, we performed immunohistochemistry on brain sections of a mouse model with amyloid deposition. The mouse model, PS1/APP, contains transgenes for both presenilin-1 with the L166P mutation and APP containing the Swedish mutation (KM670/671NL) [46]. These transgenes are under the control of the Thy1 promoter, thereby limiting expression primarily to brain tissue. Both 1A8 and 7C8 stain plaques in PS1/APP, comparable to a total Aβ antibody (Fig 6). No specific staining was observed in non-transgenic mice or with an isotype control antibody (Fig S6).

**Figure 6.**
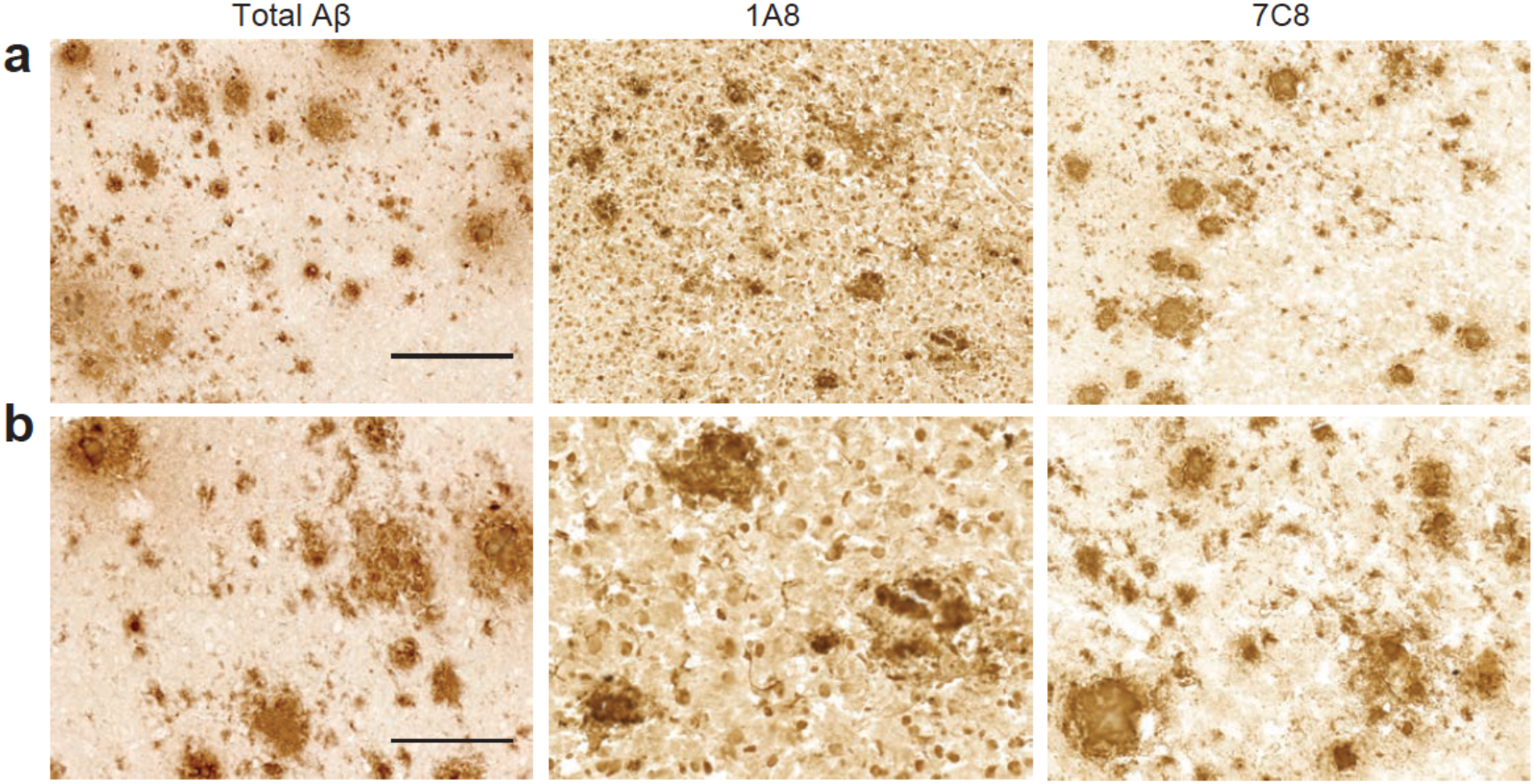
1A8 and 7C8 target reactivity in PS1/APP mouse brain. The target reactivity of 1A8 and 7C8 was evaluated in tissue slices from PS1/APP mouse brain. 12μm thick sections were post-fixed and stained with 1A8, 7C8, a positive control anti-Aβ antibody, at 10μg/ml. The signal was detected with horseradish peroxidase (HRP)-based detection. (A) A larger field of view shows the density of plaque staining in the region. (scale bar = 250 μm) (B) A higher magnification shows dense and diffuse plaque staining. (scale bar = 100 μm)

### 1A8 and 7C8 staining of human AD frontal cortex

Immunohistochemical studies were next conducted on human AD brain parenchyma. Because of the N-terminal, yet conformation-sensitive activity of 1A8 and 7C8, we analyzed the staining in adjacent serial sections of frontal cortex, and compared it with the anti-Aβ N-terminus 6E10 antibody (Fig 7A). The staining pattern showed by 1A8 and 7C8 revealed increased detection of dense neuritic plaques and large diffuse plaques within the same regions, compared with the positive control antibody (Fig 7B). Intraneuronal staining was observed with both 1A8 and 7C8, and was particularly prevalent with 1A8. Finally, parenchyma vessels were identified, revealing increased staining pattern around the vessels with both 1A8 and 7C8 (Fig 7C).

**Figure 7.**
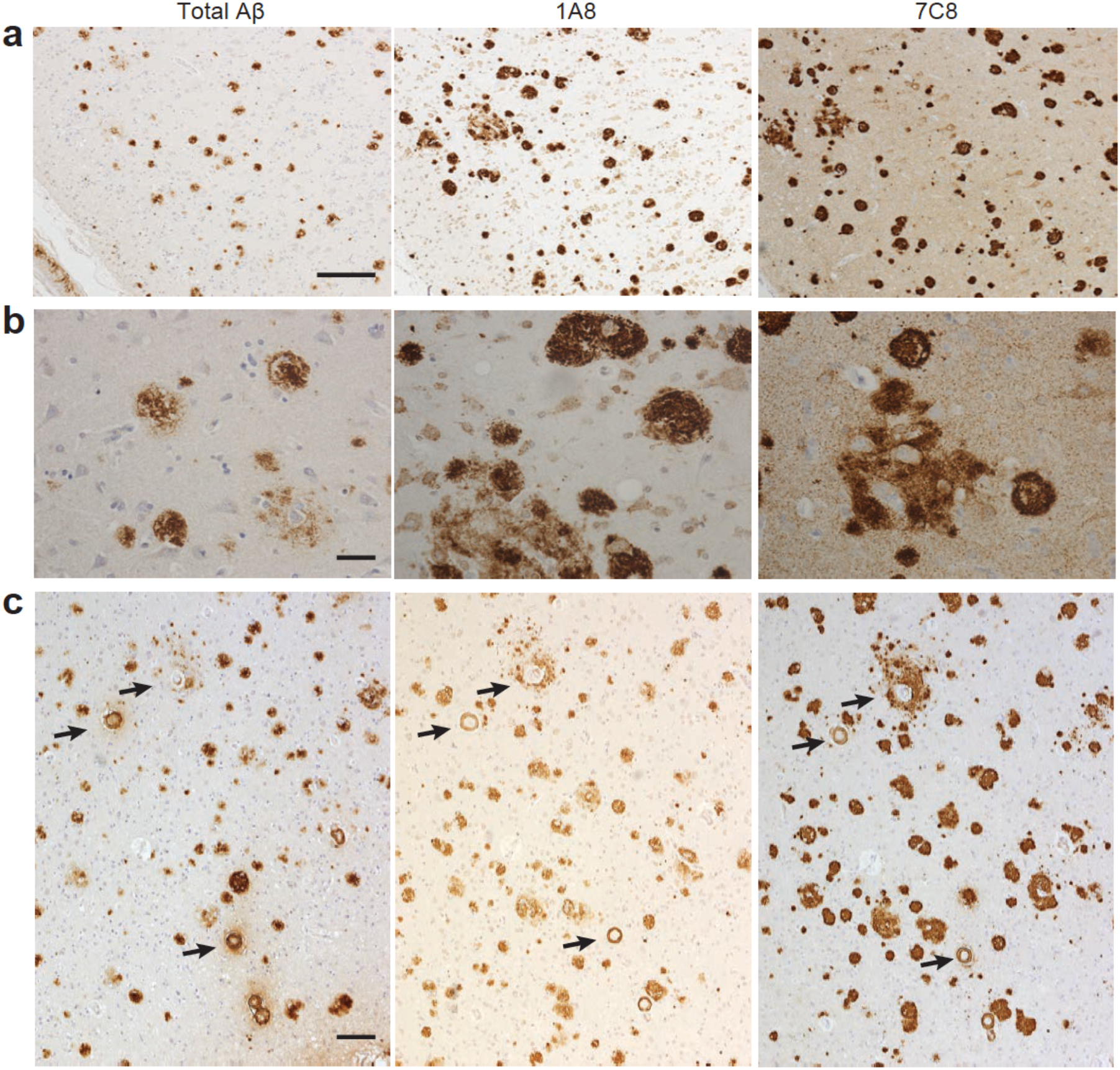
1A8 and 7C8 target reactivity in human AD brains. The target reactivity of 1A8 and 7C8 was evaluated in tissue slices from the frontal cortex of a patient diagnosed with AD (Braak V-VI) and cerebral amyloid angiopathy. 5μm thick consecutive formalin-fixed paraffin-embedded (FFPE) sections were stained with the positive control anti-Aβ antibody, 6E10 (1:1000), 1A8 (0.05μg/ml) or 7C8 (0.05μg/ml). The signal was detected using an HRP-DAB based detection system and the nuclei counterstained with Mayer’s hematoxylin. (A) A large field of view of serial sections shows dense and diffuse plaque staining with 1A8 or 7C8. (scale bar = 400μm) (B) A higher magnification shows intracellular staining with 1A8 and 7C8 that is absent with 6E10. (scale bar = 50 μm) (C) A large field of view shows staining around parenchymal vessels (arrows) in the frontal cortex. (scale bar = 100 μm)

## Discussion

Therapeutic application of conformation-selective anti-amyloid antibodies that target the pathophysiological forms of the soluble amyloid intermediate is a compelling approach to disease mitigation. Although an unproven strategy to address the huge unmet need for disease-modifying therapeutics for AD, there are four biologics in clinical trials that have reactivity against aggregated forms of Aβ. Developing conformation-selective antibodies to the extremely transient intermediate in amyloid aggregation has been a challenge. The present study describes the use of a novel CLABP, mtNUCB1, which stabilizes soluble Aβ42 protofibrillar structures [32] for use as immunogen for the discovery of conformation-sensitive, anti-Aβ protofibril mAbs (Fig 1). We hypothesized that the mtNUCB1-capped Aβ42 protofibrils would provide neoepitopes that retain protofibril structural characteristics.

mtNUCB1-capped Aβ42 protofibrils were enriched using SEC and tested with ELISA, AFM and immunoEM (S1 Fig). The results recapitulate our previous observations that mtNUCB1-Aβ protofibrils constitute a stable complex [32]. The campaign using the *mt*NUCB1-Aβ protofibril as an immunogen in mice yielded a range of activity measured from the supernatants of isolated cell populations (Fig 2A). The results of the primary screening indicate that the supernatant fractions had reactivity against Aβ protofibrils alone, mtNUCB1 alone, and in some cases, mixed reactivity against both protofibrils and *mt*NUCB1 (Fig 2B). We chose 57 hybridomas with the highest reactivity to Aβ42 protofibrils for subsequent secondary screening by single-point ELISA to assess binding to Aβ42 monomers, protofibrils, mtNUCB1-Aβ42 protofibrils, and mtNUCB1 (Fig 2B). Based on their binding profile, 24 clones (showed in Fig 2B, lower panel) were selected to measure Aβ protofibril-selective binding by the label-free Octet assay (S2 Fig A) and to determine the affinity to the peptide N-terminus by SPR (S2 Fig B).

Based on the knowledge that aducanumab is a high-affinity, conformation-selective anti-Aβ protofibril mAb that has reactivity against residues 3-6 on Aβ [47, 48], we selected from the secondary screen five antibodies that showed N-terminus binding: two antibodies with the lowest and highest binding signal, 1A8 and 7C8, respectively, and three antibodies with an intermediate binding signal (1D9, 1E7, and 6G2). These five antibodies were sequenced, subcloned and re-tested for their conformation-selectivity by sandwich ELISA. The clones showed a similar profile with moderate activity against the monomeric peptide and higher binding to Aβ42 protofibrils (S3 Fig). Therefore, among those five N-terminal mAbs with preferential binding to protofibrils, we ultimately selected the two mAbs with lowest and highest N-terminal activity, 1A8 and 7C8, and thoroughly characterized them in different assays.

First, in order to evaluate 1A8 and 7C8 conformation-sensitive activity, we tested their binding to Aβ monomers and protofibrils in a sandwich ELISA assay (Fig 3A, B), a solution competition ELISA (Fig 3C, D), and with SPR (Fig 3E-H). Our results indicate that, despite binding to the N-terminal Aβ peptide, both mAbs preferentially bind to protofibrils over monomers. The solution competition ELISA displays comparable results to that reported for aducanumab [48]. When conformation reactivity is measured by SPR, the mAbs-monomer binding is followed by an almost complete dissociation curve (Fig 3E, F), but the mAbs-protofibril association seems to be extremely strong and resistant to spontaneous dissociation (Fig 3G, H). These results could at least be partially explained by the observation that the Aβ N-terminus is a disordered extension from the aggregate core [49, 50] and thus binding activity could yield an avidity effect. Furthermore, the association curve produced by 1A8 binding to monomers is reduced compared to 7C8, in line with a lower 1A8 N-terminal affinity, as previously shown in Supplementary Fig 2B.

To determine if the binding of these mAbs had a functional effect on Aβ aggregation *in vitro,* we performed a kinetic ThT aggregation assay. We observed that, while 7C8 completely inhibited equimolar concentrations of Aβ42, 1A8 only exhibited a mild effect as can be observed by a delay in the aggregation curve (Fig 4A). Interestingly, when the experiment employed Fab fragments digested from 1A8, we observed a loss of inhibitory activity (Fig 4B), suggesting that the inhibition of aggregation expressed by the 1A8 full IgG molecule is likely due to a combined two-site binding effect lacking strong N-terminal binding.

On the other hand, 7C8 Fab fragments retained the mAb inhibitory activity as indicated by a significant and concentration-dependent inhibition of aggregation (Fig 4C *vs* 4A). The analysis of the rate kinetics for Aβ aggregation in the presence of increasing concentrations of 7C8 Fab fragments indicates that the inhibitory effect is exerted through a dual mechanism by interference with both the primary and the secondary nucleation (Fig 4D). This observation is in line with the hypothesis that 7C8 can bind structural elements required for secondary nucleation inhibition, but also binds the N-terminus of Aβ, thereby inhibiting primary nucleation events.

Altogether, these data are consistent with the N-terminal binding profile reported in Supplemental Fig 2B, with 1A8 and 7C8 showing low and high N-terminal binding signal, respectively. In fact, the N-terminus activity expressed by 7C8 (S2 Fig B) combined with the other results indicates that this mAb likely binds to both monomers and protofibrils, therefore inhibiting the early stages of the amyloid aggregation and resulting, in turn, in potent aggregation inhibition (Fig 4A, C). 1A8, on the other hand, displays low binding affinity to the Aβ N-terminus (S2 Fig B) and yet conformation-sensitive anti-protofibril activity (Fig 3A, C, E, G). Thus, the more mild inhibition of Aβ42 aggregation by the full 1A8 antibody (Fig 4A) is primarily exerted through aggregate binding with an avidity effect. The avidity feature of 1A8 to the complex, multi-repeating epitopes on the amyloid aggregate highlights structural binding.

The ideal therapeutic mAb would reduce Aβ toxicity even in the absence of aggregate degradation or cellular clearance by microglial cells. We therefore employed a cellular assay of Aβ toxicity using either Aβ42 monomers or protofibrils as the starting material in a toxic insult. 1A8 and 7C8 showed robust cyto-protection with EC50 ranges below 10 nM or below 2 nM for either monomer or protofibril insults, respectively (Fig 5). These results highlight the binding effect of 1A8 and 7C8 on Aβ aggregates.

To visualize target binding *ex vivo,* we used immunohistochemistry on PS1/APP mouse brain and AD human frontal cortex. The results show that both antibodies display plaque staining of both dense and diffuse phenotypes (Fig 6 and Fig 7). In serial sections of human AD brain, the number of diffuse and dense plaques were greater than that detected with the positive control, Aβ N-terminus 6E10 antibody. Additionally, both mAbs showed moderate intraneuronal staining, whereas this pattern was largely absent with 6E10 staining (Fig 7). The observed 1A8 and 7C8 staining pattern was not due to reactivity with APP, since it differed with that of 6E10, an antibody known to recognize APP as well as Aβ. Furthermore, parenchymal vessels were stained with both antibodies and diffuse staining around vessels could be observed. Therefore, these mAbs may be useful tools for studying intraneuronal Aβ protofibril accumulation or aggregate accumulation at parenchymal vessels.

## Conclusions

We report the use of a novel protofibril immunogen that has been stabilized by the CLABP mtNUCB1. The immunization campaign using this pan-amyloid, protofibril-capping CLABP in complex with Aβ42 protofibrils yielded at least two high-affinity, conformation-sensitive mAbs, 1A8 and 7C8, that preferentially bind the Aβ42 protofibril over the Aβ monomer, though with apparent varying binding modes. Because mtNUCB1 stabilizes protofibrils from multiple amyloid sources, this approach represents a platform technology for discovering anti-protofibril antibodies that are potentially diagnostic or therapeutic for many amyloidosis syndromes.

## Acknowledgements

Funding for this project was provided through the Pfizer Centers for Therapeutic Innovation program, the Robertson Therapeutic Development Fund at Rockefeller University, the Eleanor Schwartz Charitable Trust, the Nicholson Exchange Program and the Swedish Brain Power at the Karolinska Institutet. Human brain tissue was kindly provided by the Brain Bank at Karolinska Institutet, which receives financial support from the Stockholm County Council (Core facility funding), StratNeuro Karolinska Institutet and Swedish Brain Power. We thank the Electron Microscopy Resource Facility and the Molecular Cytology Core Facility at Memorial Sloan-Kettering Cancer Center. We thank Dr. Carolina Adura of the Rockefeller University High-Throughput and Spectroscopy Resource Center for guidance and data interpretation in the use of the ProteOn XPR36 Protein Interaction Array system. We also thank Dr. Fergus Byrne for early contributions to the project and Prof. Bengt Winblad for advice and encouragement throughout.

## Authors’ contributions

WVG, ABO, PDW, and TPS designed the study. WVG and ABO prepared and characterized the immunogen, and conducted the experiments with the purified antibodies. RA, RK, KK, and KM supervised the immunization campaign and carried out the primary and secondary screening. JD performed the immunohistochemical studies on animal tissue and AL, MP, and CG performed studies on human tissue. WVG and ABO wrote the paper with critical input from CG, KMW, KM, PDW, TPS and all other authors who approved the final version.

