## Supporting Information for "Discovery of conformation-sensitive anti-amyloid protofibril monoclonal antibodies using an engineered chaperone-like amyloid-binding protein"

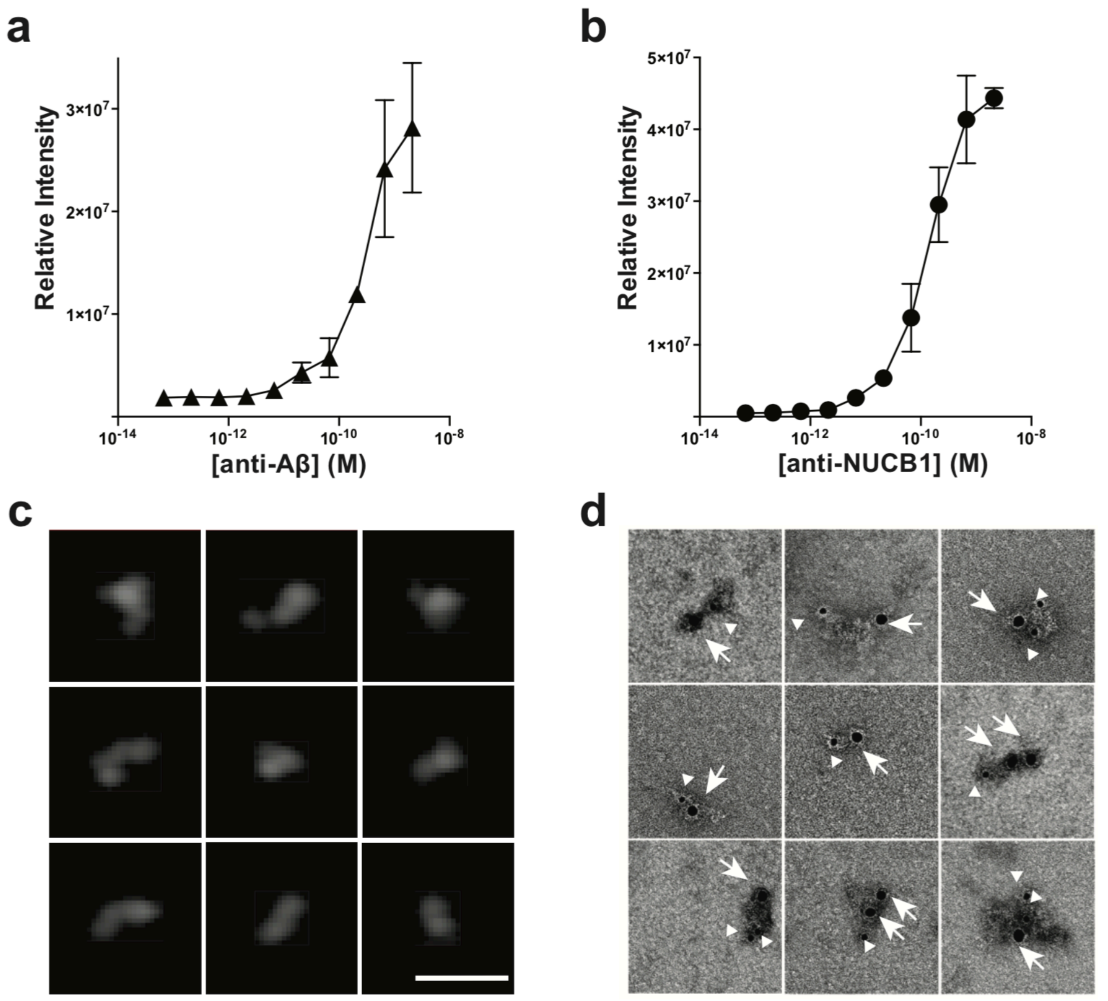


**S1 Fig. Characterization of *mt*NUCB1-stabilized Aβ-protofibril immunogen.**

*mt*NUCB1-Aβ42 protofibrils were purified by SEC and tested by direct ELISA to measure their (A) Aβ42 and (B) *mt*NUCB1 content. (C) Composite of representative protofibrils imaged by AFM (scale bar = 40nm) and (D) double immunoEM where *mt*NUCB1 and Aβ42 are bound by 12 nm or 6 nm gold particles, respectively (100 nm x 100 nm square). Arrows indicate *mt*NUCB1, arrowheads indicate Aβ42.

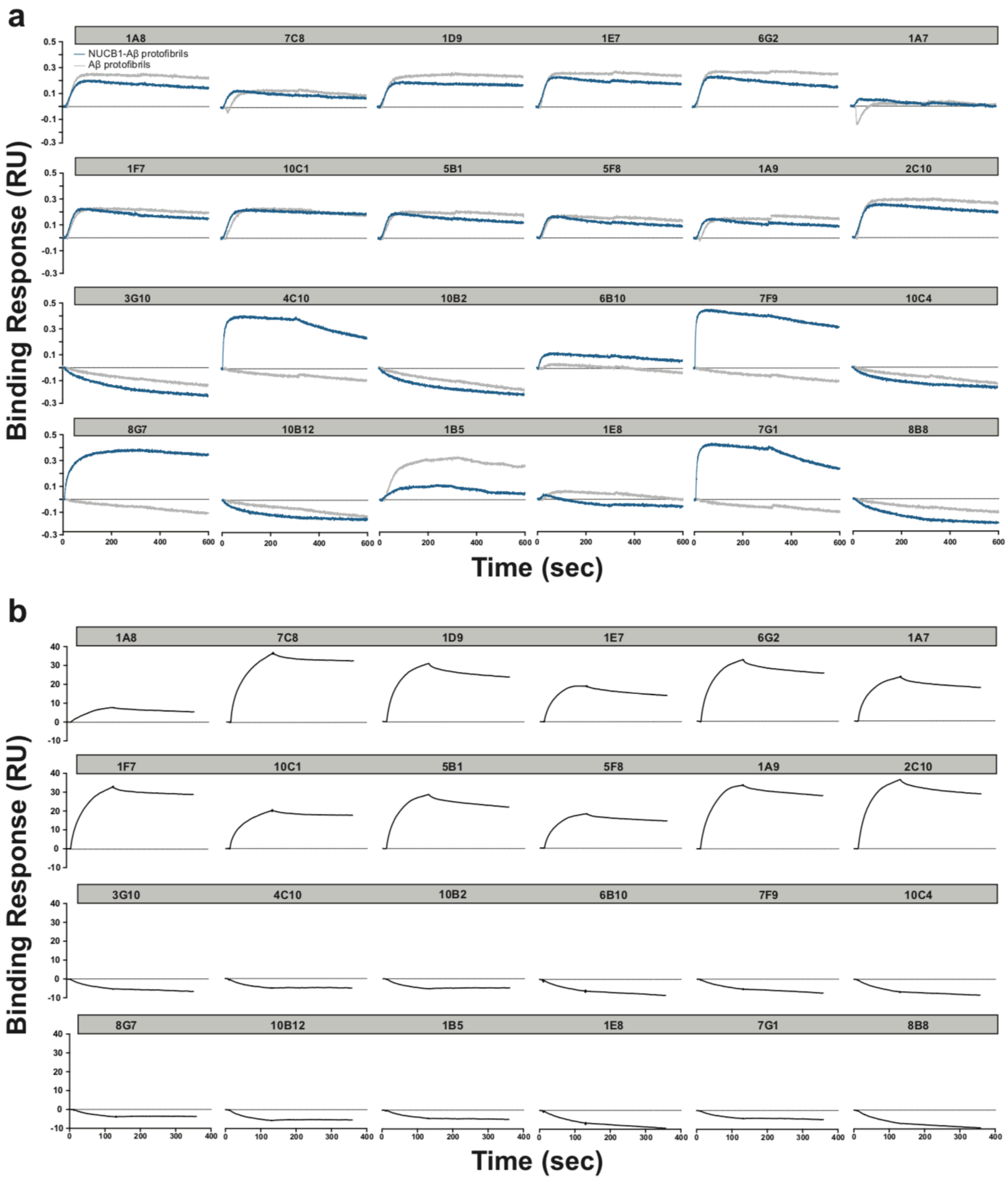


**S2 Fig. Characterization of Antibodies by Octet and Biacore.**

(A) Binding of the selected antibodies to the immunogen (*mt*NUCB1-Aβ protofibrils) and to Aβ protofibrils, measured by the label-free high-throughput Octet assay. The antibodies where captured with an anti-mouse IgG Fc biosensor and the sensorgram binding responses (nm) was measured for 600 sec. (B) Binding of the selected antibodies to the Aβ peptide DAE10-EG in the Biacore assay to determine N-terminus binding. The antibodies where captured with an anti-mouse Fc coupled biosensor and a single concentration of 500 nM DAE10-EG peptide was flowed over them. The binding responses (RU) were measured for 600 seconds.

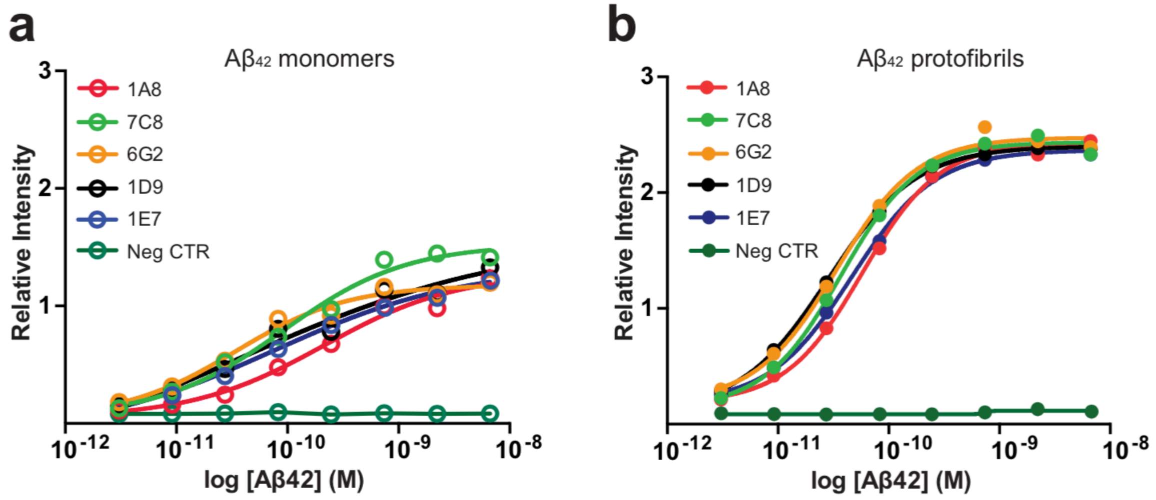


**S3 Fig. Conformation-specific binding to monomers and protofibrils.**

Based on the results of the primary and secondary screening, the heavy and light chain variable regions ($V_H$ and $V_L$) of 5 clones (1A8, 7C8, 6G2, 1D9, and 1E7) were cloned into antibody expression vectors expressing human constant regions (Fc). Antibodies were produced recombinantly, and screened at a range of concentrations for reactivity against Aβ (A) monomers and (B) protofibrils in order to determine $EC_{50}$ values. The relative intensity binding was compared to a human IgG1 isotype control (Neg CTR).

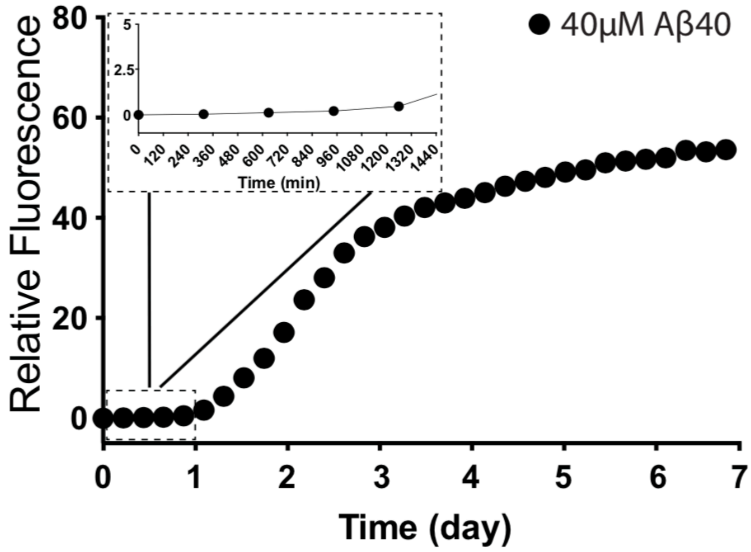


**S4 Fig. Aggregation kinetics of Aβ40.**

Aβ40 (40 *μ*M) was incubated together with thioflavin T (20 *μ*M) and incubated for 7 d at 25 °C under quiescent conditions. The insert shows the first 24h of incubation. Data are presented as mean ± SEM.

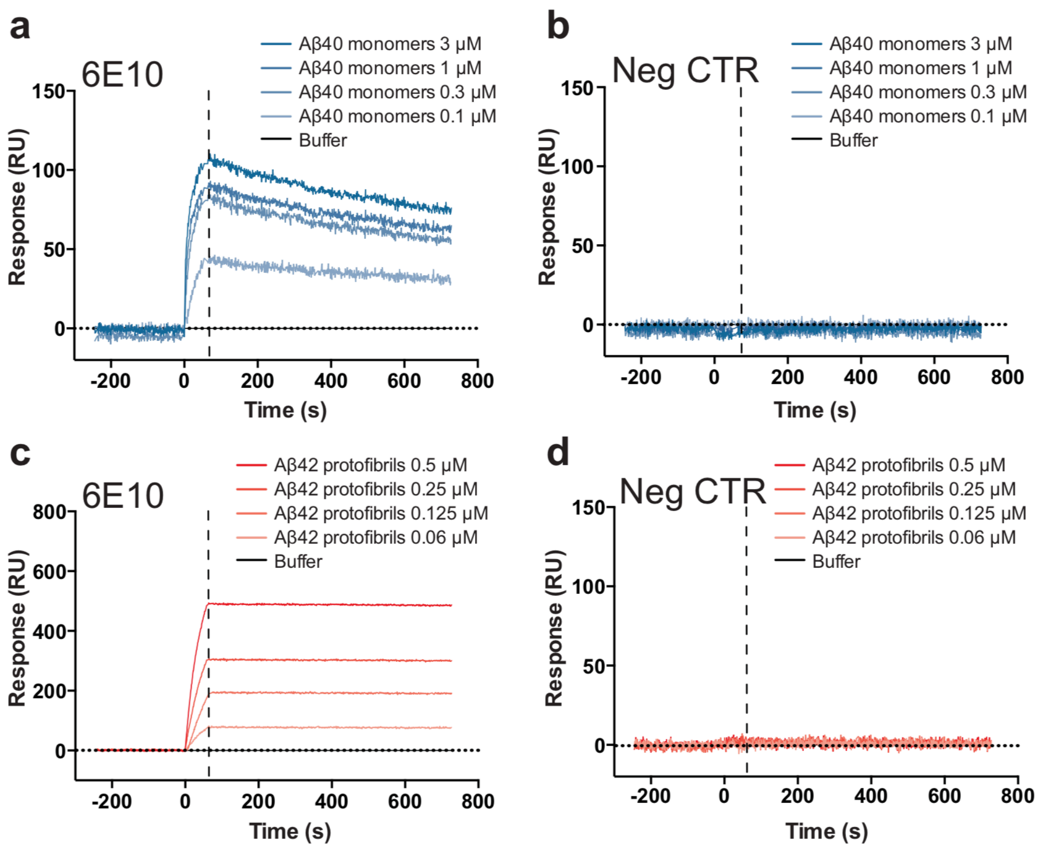


**S5 Fig. Binding of the positive and negative control antibodies to Aβ protofibrils and monomers.**

The binding of (A,B) the positive control anti-Aβ 6E10 antibody and (C,D) the negative IgG control 1D4 antibody to (A,C) Aβ40 monomers and (B,D) Aβ42 protofibrils was tested by SPR. Aβ40 monomers and Aβ42 protofibrils were prepared at a 10 $\mu$M initial concentration, diluted and flown at different concentrations (monomers: 3, 1, 0.3, and 0.1 $\mu$M; protofibrils: 0.5, 0.250, 0.125, and 0.06 $\mu$M) for 60 s over each antibody (300 nM) previously immobilized on the chip (RL = 6500). Data are normalized by interspot and buffer and presented as mean ± standard error of the mean (SEM).

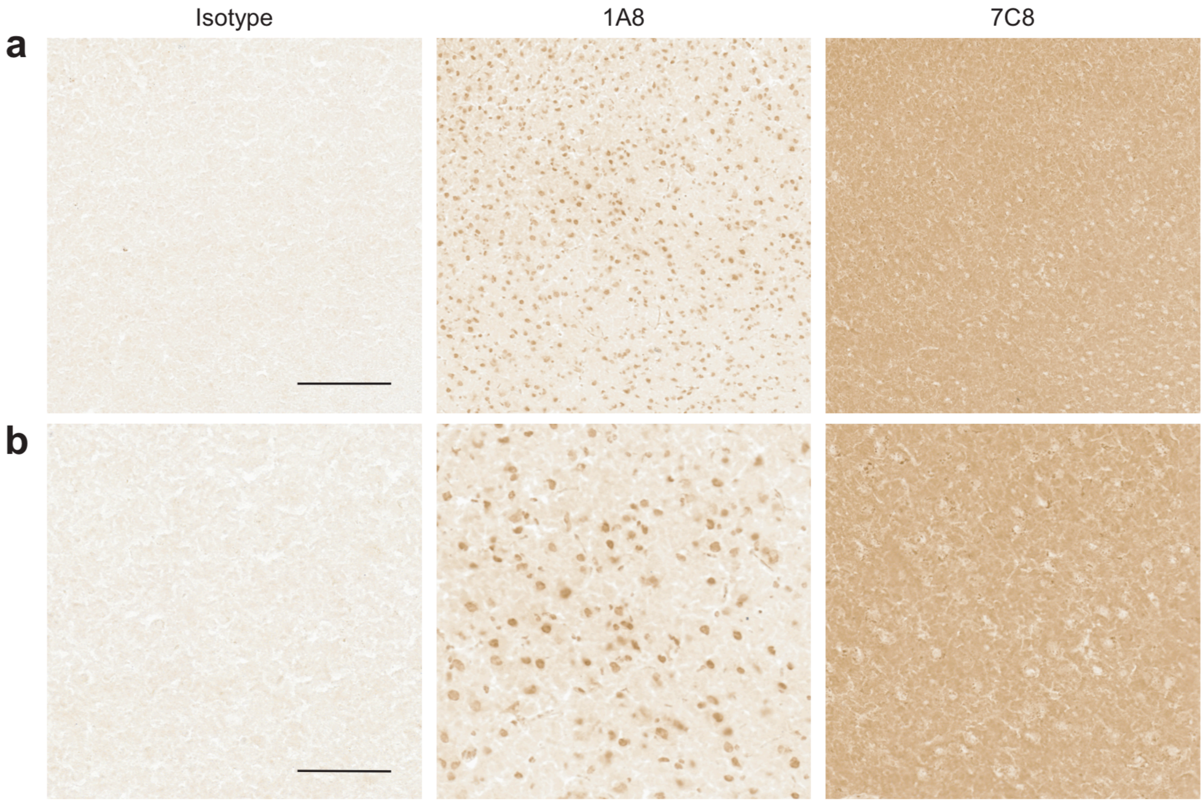


**S6 Fig Binding of 1A8 and 7C8 to non-transgenic mouse brain tissue.** The reactivity of 1A8 and 7C8 was evaluated in slices from non-transgenic control brains. 12 *μ*m thick sections were post-fixed and stained with 1A8, 7C8 or a human IgG1 isotype control antibody, at 10 *μ*g/ml. The signal was detected with horseradish peroxidase (HRP)-based detection. (A) A larger field of view (scale bar = 250 μm). (B) A higher magnification shows no staining of tissue (scale bar = 100 μm).

| | | |
|---|---|---|
| Model used: | Mulit-Step_Secondary_Nucleation_Dominated_noseed | |
| Data normalized: | yes | |
| Mean squared error: | 0.004863637 | =sum[(y_i - f(x_i))^2]/(N_datapoints - N_free_parameters) |

| Dataset<br>Units | kp*kn<br>conc^{-n_c} time^{-2} | mo | nc<br>unitless | KM<br>conc^{n_2} | n2<br>unitless | kptimesk2<br>conc^{-n_2-1} time^{-2} |
|---|---|---|---|---|---|---|
| Aβ42 10 μM + 7C8 Fab 2.5 μM | 81500000 | 1.50E-06 | 2.07 | 7.32E-12 | 1.95 | 8.91E+14 |
| Aβ42 10 μM + 7C8 Fab 5 μM | 100000000 | 1.10E-06 | 2.01 | 1.28E-11 | 1.98 | 6.00E+14 |
| Aβ42 10 μM + 7C8 Fab 7.5 μM | 100000000 | 6.20E-07 | 2 | 4.81E-11 | 2 | 6.00E+14 |
| Aβ42 10 μM + 7C8 Fab10 μM | 97300000 | 5.50E-07 | 2 | 1.15E-11 | 2 | 6.00E+14 |

**S1 Table. Kinetic analysis of 7C8 Fab fragments.**

The kinetic data of 10 μM Aβ42 aggregation in the presence of varying concentrations of 7C8 Fab fragments were analyzed with Amylofit to perform individual simulations. All the parameters [initial monomer concentration (m0), reaction order of primary nucleation (nc), and reaction order of secondary nucleation (n2)] were set to Global constant and each time one of the rate constants (primary nucleation, elongation, or secondary nucleation) was set to 'Fit' while the others were set to 'Global fit'.
